# Age-related change in transient gamma band activity during working memory maintenance through adolescence

**DOI:** 10.1101/2022.07.24.501317

**Authors:** Shane D. McKeon, Finnegan Calabro, Ryan V. Thorpe, Alethia de la Fuente, Will Foran, Ashley C. Parr, Stephanie R. Jones, Beatriz Luna

## Abstract

Adolescence is a stage of development characterized by neurodevelopmental specialization of cognitive processes. In particular, working memory continues to improve through adolescence, with increases in response accuracy and decreases in response latency continuing well into the twenties. Human electroencephalogram (EEG) studies indicate that gamma oscillations (35-65 Hz) during the working memory delay period support the maintenance of mnemonic information guiding subsequent goal-driven behavior, which decrease in power with development. Importantly, recent electrophysiological studies have shown that gamma events, more so than sustained activity, may underlie working memory maintenance during the delay period. However, developmental differences in gamma events during working memory have not been studied. Here, we used EEG in conjunction with a novel spectral event processing approach to investigate age-related differences in transient gamma band activity during a memory guided saccade (MGS) task in 164 10- to 30-year-olds. Total gamma power was found to significantly decrease through adolescence, replicating prior findings. Results from the spectral event pipeline showed age-related decreases in the mean power of gamma events and trial-by-trial power variability across both the delay period and fixation epochs of the MGS task. In addition, we found that while event number decreased with age during the fixation period, it did not appear to change during the delay period resulting in an increasing difference between the number of events during fixation and delay period with development, suggesting that as working memory develops there is greater specificity for gamma events supporting working memory. While average power of the transient gamma events was found to mediate age-related changes in total gamma power, the number of gamma events was unrelated to total power, suggesting that the power of gamma events may underlie the sustained gamma activity seen in EEG literature while the number of events may directly support age-related improvements in working memory maintenance. Our findings provide compelling new evidence for mechanistic changes in neural processing characterized by refinements in neural function as behavior becomes optimized in adulthood.

## 1. INTRODUCTION

Working memory continues to improve through adolescence into adulthood^1,2^ in parallel with brain maturation^2– 4^ and optimization of brain function^5,6^. Behavioral studies using verbal and visuospatial tasks have consistently shown that working memory accuracy and latency continue to improve well into the twenties^5–9^. Supporting this cognitive maturation, fMRI studies have found that while brain regions involved in working memory are online by childhood, there is a continued refinement and integration of specialized regions that lead to stabilized neural activity and an improvement in behavioral performance, such as latency and accuracy^5,6^. In particular, there is substantial evidence that in addition to overall improvements in working memory performance through adolescence, there are significant increases in the reliability with which individuals performed these functions, seen as a reduction in the variability of trial-to-trial accuracy and latency^5,6^.

Previous developmental electroencephalogram (EEG) studies have shown robust trends in declining power during resting state EEG across early childhood, middle childhood, and into adolescence^10^. More specifically, agerelated decreases in absolute power have been found to decrease in the slow wave (0.5 – 7.5 Hz)^11^, alpha (8 – 12 Hz), and beta (12.5 – 34.5 Hz) bands in the frontal, temporal, parietal, and occipital lobes^12,13^, as well as the theta (3.5 – 7.5 Hz)^14^, and gamma (30 – 70 Hz)^15^ bands. In terms of relative power, there have been age-related increases in the alpha^16,17^, and beta^16^ bands, as well as a redistribution from lower to higher frequency bands with posterior regions reaching adult levels of power before frontal regions^18^.

Human EEG and animal electrophysiological data have identified that specific aspects of these neural signals, including gamma oscillations, support working memory processes^15,19^, particularly in the dorsolateral prefrontal cortex (DLPFC)^6,20^ where gamma power during working memory tasks was increased compared to baseline^19^. Historically, these gamma oscillations have been characterized as reflecting sustained activity during the maintenance epoch of working memory tasks in both non-human primate and human studies^21–25^. However, recent human and non-human primate studies have shown that at the trial level, neural activity occurs in burst-like events, defined as transient events of neural activity where not only are the amplitude and frequency important, but the timing, duration, and rate of the events may play a key role in supporting higher order cognitive functions^21,26–30^. Working memory delay activity has since been shown to be non-stationary, with spiking occurring sporadically^28,30,31^. Between these active states, working memories may be stored as temporary changes in synaptic weights through gamma band associated spiking^23,28,30,32^. However, while EEG provides a means for assessing working memory-dependent gamma events, no studies to our knowledge have yet characterized in developmental populations the trial-level spectral events associated with the maturation of working memory. Here, we build off previous EEG studies using established averaging approaches of sustained activity by investigating whether non-stationary working memory activity can provide insight into the underlying mechanisms of working memory during adolescent development. We apply a novel EEG analysis pipeline^33^ to define the developmental trajectory of gamma band events at the trial level, their association with working memory throughout adolescence into young adulthood, and their relationship with traditional EEG analyses of sustained activity. These results demonstrate aspects of the spectral events themselves which account for prior findings of decreased overall gamma power, while highlighting novel developmental changes in the rate of task-dependent gamma events which may constitute a distinct developmentally sensitive process supporting the maturation of working memory maintenance.

## 2. METHODS

### 2.1 Participants

One hundred and sixty-four participants (87 assigned female at birth), between 10 and 32 years of age participated in this study. Participants were recruited from the community and were excluded if they had a history of head injury with loss of consciousness, a non-correctable vision problem, a history of substance abuse, a learning disability, or a history of major psychiatric or neurologic conditions in themselves or a first-degree relative. Participants were also excluded if they reported any MRI contraindications, such as non-removable metal in their body given other neuroimaging aspects of the broader project. Participants or the parents of minors gave informed consent with those under 18 years of age providing assent. Participants received payment for their participation. All experimental procedures were approved by the University of Pittsburgh Institutional Review Board and complied with the Code of Ethics of the World Medical Association (Declaration of Helsinki, 1964).

### 2.2 Memory Guided Saccade Task

Participants performed a memory guided saccade (MGS) task to assess working memory (see Figure 1). The trial began with fixation to a blue cross for 1 sec. The participant was then presented with a peripheral cue in an unknown location along the horizontal midline (12.5 or 22.2 degrees from central fixation to left or right of center), where they performed a visually guided saccade (VGS) to the target and maintained fixation. Once the cue disappeared, the participant returned their gaze to the central fixation point and fixated for a variable delay epoch (6-10 sec) during which they were to maintain the location of the peripheral target in working memory. Once the central fixation disappeared the participant performed a memory guided saccade to the recalled location of the previous target. The trial ended when participants were presented with a white fixation cross that served as the ITI (1.5-15sec). Participants performed 3 runs of the MGS task, each containing 20 trials.

**Figure 1.**
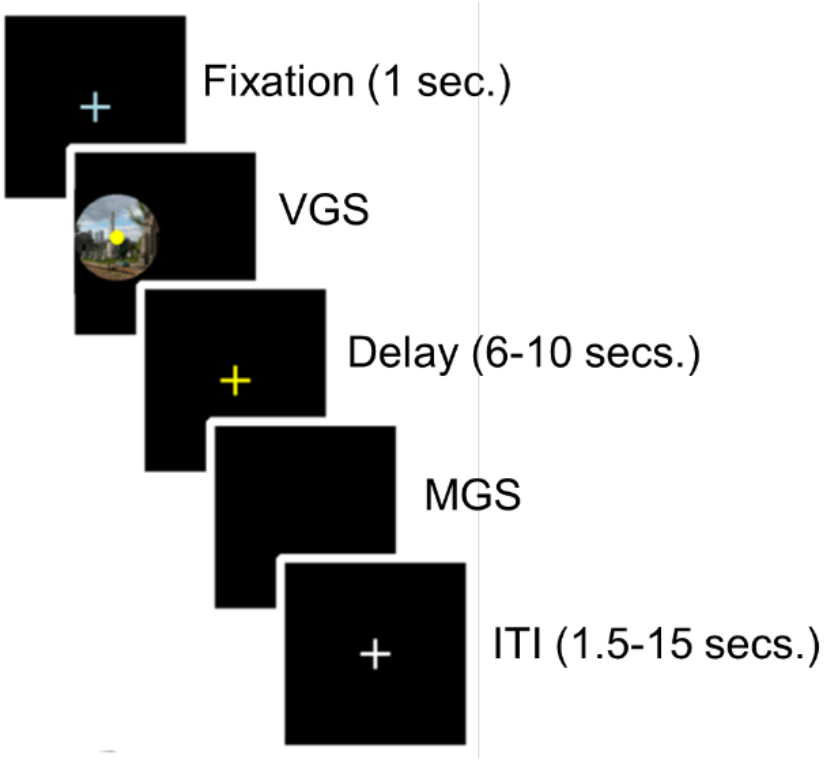
Memory Guided Saccade Task. Epochs from top to bottom: Participants were presented with a fixation cross. Once fixation was extinguished, participants performed a saccade to a peripheral dot stimulus on top of a scene (the scene is intended for an incidental memory task for a separate aim of this project). After the peripheral target was extinguished, participants were presented with a yellow fixation cross while remembering the location of the previous target. When the fixation cross was extinguished, participants performed a second saccade to the remembered target location. A variable ITI (1.5-15 sec) with a fixation white cross occurred between trials.

Task performance was assessed based on horizontal electrooculogram (hEOG) channels recorded from facial muscles (see acquisition details below). At the start of the session, participants performed a calibration procedure in which they fixated a series of 20 dots sequentially, positioned along the horizontal midline and spanning the width of the screen. These were used to generate a calibration curve relating hEOG voltage to horizontal screen position. Eye position during the MGS task was used to derive output measures using this calibration data by aligning hEOG signals to different task triggers. These were used to calculate VGS & MGS response latencies, as the time difference between the beginning of the recall epoch and the initiation of the VGS and MGS eye movements respectively, and saccadic accuracy, measured as the closest fixation point during the recall period to the fixated location during the initial visually guided fixation epoch.

### 2.3 Electrophysiological Data Acquisition and Preprocessing

Concurrent EOG and high-impedance EEG was recorded with a Biosemi ActiveTwo with Actiview software (Minimitter, Bend, OR) 64-channel EEG system, in accordance with the international 10-20 system^33^, located in an electromagnetically shielded room while participants performed the MGS task (described above). The task stimuli were presented on a computer 80 cm from the subject, and head position was maintained using a chin rest. Initial data were sampled at 512 Hz and down sampled to 150 Hz during preprocessing. Data were referenced to external electrodes corresponding to the mastoids due to their proximity to the scalp and low signal recording. An initial bandpass filter was set to 0.5-75 Hz. We used a revised version of an established protocol (https://sccn.ucsd.edu/wiki/Makoto's_preprocessing_pipeline, retrieved April 23, 2020) for preprocessing compatible with EEGLAB^34^. This protocol removes flatline channels (maximum tolerated flatline duration: 8 seconds), low-frequency drifts, noisy channels (defined as more than 5 standard deviations from the average channel signal), large amplitude artifacts, and incomplete segments of data^35,36^. Deleted channels were replaced with interpolated data from surrounding electrodes. Continuous signals were divided into epochs, time-locked to stimulus onset (∼1-3s). Data epochs were cleaned with an amplitude threshold of -500 to 500μV to exclude eye blinks. The resulting data were referenced to the average of all electrodes. As a final preprocessing step, independent component analysis (ICA) was performed to identify eye-blink artifacts and remove their contribution to the data.

### 2.4 Power Analysis

Spectral power was computed for every electrode from the 3000 – 4000ms window of the delay epoch of the task to avoid artifact from preceding eye movements and preparation from an imminent response, and from the 1 second inter-trial fixation epoch. Power spectra was derived from a fast Fourier transform (FFT). Power (µV^2^) was then calculated for the gamma frequency band (35-65 Hz). Power from all 6 electrodes representing the left and right DLPFCs (F3, F4, F5, F6, F7, and F8) were averaged together. The resulting measure is referred to as “total” gamma band power throughout the manuscript.

### 2.5 Spectral Analysis

Spectral data were computed for every electrode from the 3000 – 4000ms window of the delay epoch of the task to avoid artifact from preceding eye movements and preparation from an imminent response, and from the 1 second inter-trial fixation epoch. The spectrograms of the data were calculated from 20 to 70 Hz by convolving the signals with a complex Morlet wavelet^37^ of the form

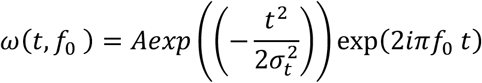

for each frequency of interest *f*_0_, where *σ* = *m*/(2*πf*_0_). The normalization factor was 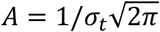 and the constant *m*, defining the compromise between time and frequency resolution, was 7, consistent with previous literature^27^. Time-frequency representations of power (TFR) were calculated as the squared magnitude of the complex wavelet-convolved data. The TFR was normalized to the median power value for each frequency band, derived from all the power values within the identified stimulus windows of the delay and fixation epochs. Normalized TFR values were calculated in factors of median (FOM) for each frequency, separately for each participant/trial.

### 2.6 Region of Interest Selection

Average gamma power (µV^2^) was first calculated by averaging the time-frequency representations of power (TFR) across time and frequency for all participants. Analyses were limited to electrodes that had well defined activity in the gamma band of adult participants during the MGS task. To accomplish this, we identified electrodes with gamma activity present by computing total power from the time- and frequency-band averaged TFR across adult subjects (18+ years of age) for each electrode. Power values were plotted across brain topography to highlight regions of increased power. Using this information, coupled with previous knowledge of regions involved with working memory^6,20^, electrodes corresponding to the right (F4, F6, and F8) and left (F3, F5, and F7) DLPFC were selected for further analysis (see Results, and Fig 4). To verify the presence of gamma band activity, spectrograms for each electrode were calculated from 20 to 70 Hz.

### 2.7 Defining Gamma Events and Features

Gamma events were defined as local maxima in the trial-by-trial TFR matrix for each frequency value at the maxima that fell within the gamma band (35 – 65 Hz) and the power exceeded a cutoff of 6x the median power^27^. Local maxima were found using the Matlab function ‘imregionalmax’. Custom software for identifying transient events and event features is written in Matlab and available at https://github.com/jonescompneurolab/SpectralEvents. Trial average and trial-by-trial variability of each feature were calculated by using the average and standard deviation of each measure (event power, number, and duration) across trials for the delay and fixation epochs separately.

#### Gamma Event Power

Gamma event power was defined as the normalized TFR value (units of FOM) at the local maximum that defines each event. The trial mean event power was defined as the power of all events averaged in the 3000 – 4000 ms window of the delay epoch and the 0 – 1000 ms window of the fixation epoch.

#### Gamma Event Number

Gamma event number was calculated as the number of gamma events in the 3000 – 4000 ms window of the delay epoch and the 0 – 1000 ms window of the fixation epoch.

#### Gamma Event Duration and Frequency Span

Event duration and frequency span are calculated as full-width-half-maximum (FWHM) from the event maxima in the time and frequency domain, respectively.

All spectral measures (power, number of events, duration of events, and variability of each) were averaged across electrodes F3, F5, and F7 to create average spectral event measures for the left DLPFC, and F4, F6, and F8 for the right DLPFC based on the region of interest analysis performed on the total TFR power of each epoch window.

### 2.8 Statistical Analysis

To examine age-related effects in behavioral measures, including accuracy and response latency, generalized additive models (GAMs) were implemented using the R package mgcv^38^. Preliminary outlier detection was conducted on a trial level basis. Express saccades of less than 100ms, believed to be primarily driven by subcortical systems^39^, were excluded. Position error measures greater than 23 degrees from the target were excluded as implausible since they exceeded the width of the screen. The remaining trials for each participant were combined, creating average performance measures and trial-by-trial variability measures for each participant. Finally, for group level analyses, outlier detection was performed, excluding participants more than +/-2 SDs away from the mean. A separate model was performed for each of the behavioral measurements: MGS accuracy, MGS latency, and VGS latency, as well as the trial-by-trial variability of each measure. To correct for multiple comparisons between the six behavioral measures, Bonferroni correction was employed which resulted in a corrected alpha of .008 (*p* = .05/6 = .008). For all behavioral measures showing significant age-related effects, we performed analyses to identify specific periods of significant age-related differences. To do so, a posterior simulation was performed on the first derivative of the GAM model fits. Consistent with previous work^38,40,41^, 10000 simulated GAM fits and their derivatives (generated at age intervals of 0.1 years) were computed from a multivariate normal distribution; vector means and covariance of which corresponded to the fitted GAM parameters. Confidence intervals (95%) were generated from the resulting derivatives. The periods of age-related growth were derived from the ages corresponding to when the confidence interval did not include zero (p < 0.05).

We used a similar GAM approach to assess age-related trends in the average power derived from the FFT and for gamma band spectral event properties. In each, we first tested for age by epoch interactions and if the interaction term was not significant, it was removed from the model and epoch was included as a covariate. Values greater than +/-2 SDs away from the variable mean were excluded. Next, a separate model was performed for each of the spectral event measurements: power, duration, number of events per trial, the trial-by-trial variability of each measure. We first tested for age by hemisphere and age by epoch interactions on each measure, and in the event that the interaction term was not significant, it was removed from the model and hemisphere and epoch were included as covariates. Trials greater than +/-2 SDs away from the variable mean were excluded. To correct for the six spectral event measures, a corrected p value of 0.008 was used.

To assess relationships between the significant age-related EEG activity and behavior, each delay epoch spectral measure in which we found significant age effects was compared with the memory guided saccade behavioral measures (accuracy, measured in degrees from the correct target location, and response latency), again using GAMs. To correct for the resulting six comparisons, a Bonferroni-corrected p value of 0.008 was used as the criterion for significance.

To test for relationships between the age-related changes in the spectral event measures and agerelated changes in total power, mediation analyses were implemented using the R package *mediation*^42^. Unstandardized indirect effects were computed for each of 1,000 bootstrapped samples, and the 95% confidence interval was computed by determining the indirect effects at the 2.5^th^ and 97.5^th^ percentiles.

## 3. RESULTS

### 3.1 Behavioral Performance

Consistent with our prior findings using the MGS task, behavioral performance improved with age for all MGS metrics including increased accuracy (*F* = 40.89, *p* = 1.2e-15; Figure 2A), decreased response latency (*F* = 21.25, *p* = 0.01; Figure 2B), and decreased trial-to-trial variability in both accuracy (*F* = 48.93, *p* = 8.82e-07; Figure 2C) and response latency (*F* = 61.48, *p* = 1.2e-15; Figure 2D). Significant developmental improvements were found to occur throughout adolescence (11-22 years of age) for MGS accuracy and between 11-21 years of age for trial-by-trial variability for MGS accuracy (Figure 2A and 2C). MGS latency was found to have significant growth rates in early adolescence (11-13 years of age, Figure 2B), while decreases in MGS variability in latency continued into the twenties (Figure 2D).

**Figure 2.**
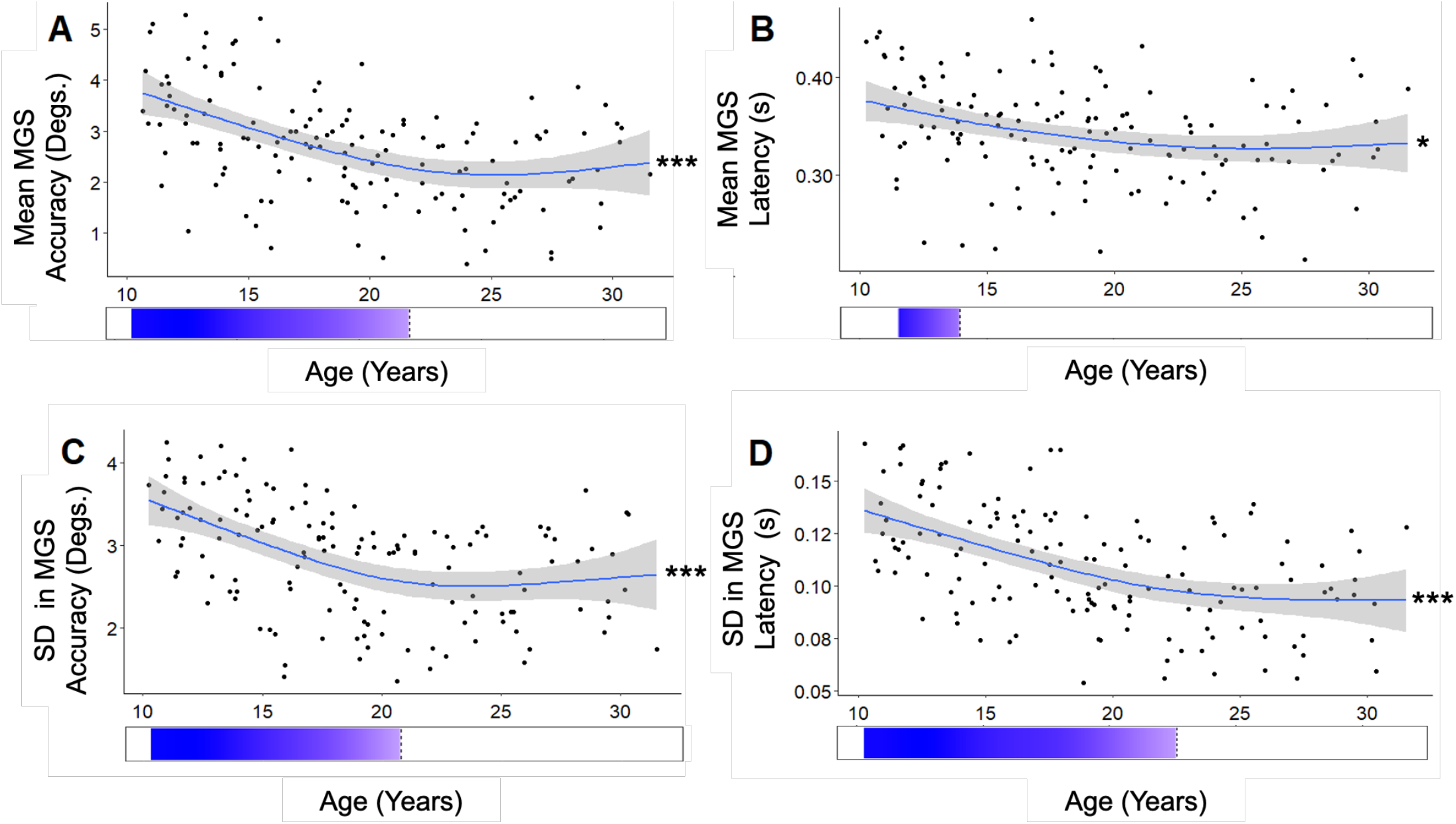
**A**. Mean MGS accuracy (degs.). Growth rates show significant rates of improvement between 11-22 years of age **B**. Mean MGS response latency (s). Growth rates show significant rates of improvement between 11-13 years of age **C**. Trial-by-trial variability in MGS accuracy (degs.). Growth rates show significant rates of improvement between 11-21 years of age **D**. Trial-by-trial variability of MGS response latency. Growth rates for show significant rates of improvement between 11-23 years of age. Blue bars indicate regions of significant agerelated improvement in MGS performance based on the derivative of the GAM model fit. (* p < 0.01; ** p < 0.001; *** p < 0.0001)

### 3.2 Gamma Power Present in Left and Right DLPFC

Given our hypothesis based on prior literature that gamma power would decrease with age, we first identified the presence of gamma band power in adults (age range 18 – 32, N = 87) to characterize the extent of activity in a mature cohort. Average gamma power was calculated across our adult cohort and plotted across brain topography to select regions of interest for investigating age-related differences in transient gamma events during the working memory delay and fixation epochs. Figure 3 depicts the presence of gamma power across all electrodes of adults during the 3000-4000ms time window of the delay epoch, with hotspots identified in bilateral DLPFC. Based on this result, and coupled with prior work of neural signals in the DLPFC supporting working memory^9,30^, bilateral DLPFC regions (right: F4, F6 and F8, left: F3, F5, and F7) were selected for further analysis. Given apparent differences in the magnitude of gamma power between left and right DLPFC, in subsequent statistical analyses we include terms for hemisphere and age by hemisphere interactions.

**Figure 3.**
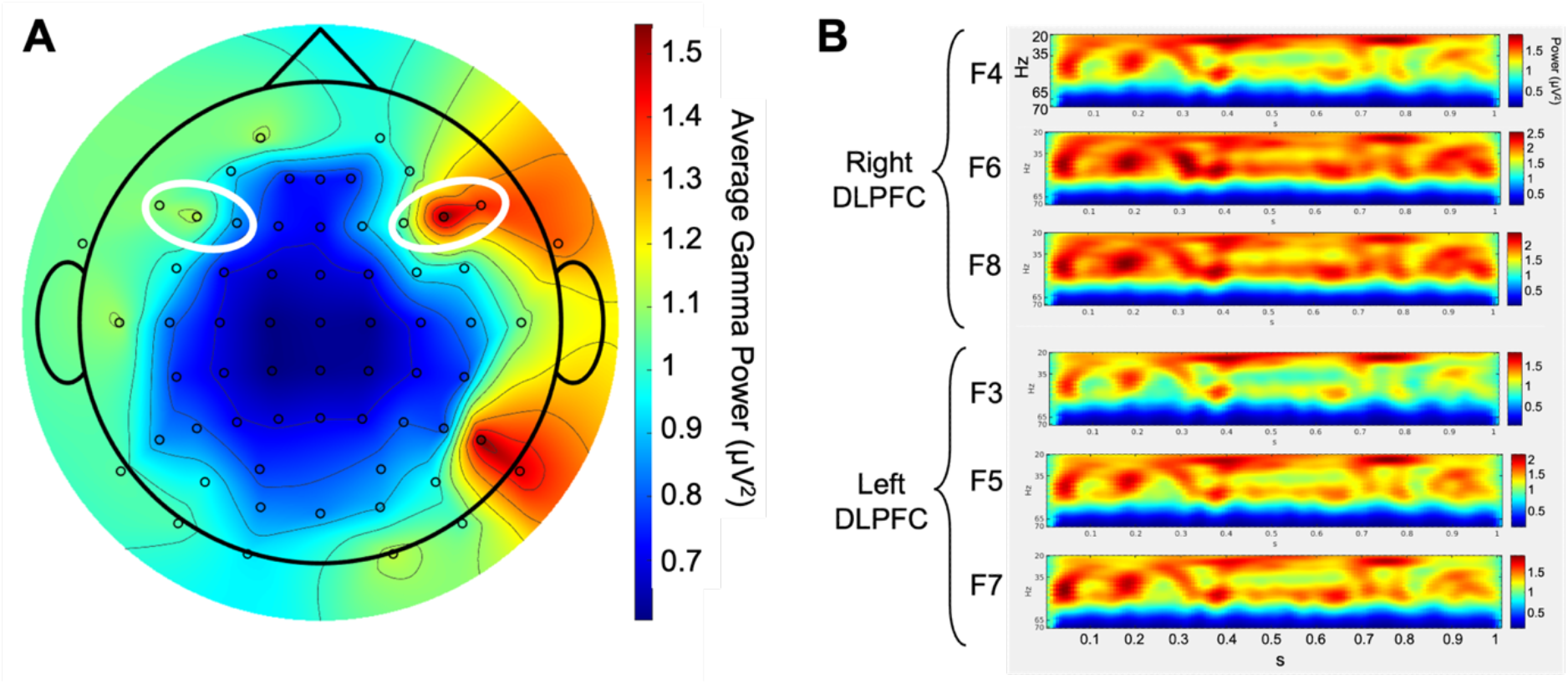
**A**. Average gamma power (µV^2^) across the entire adult cohort (18+ years of age), plotted across the brain topography. The white circles denote the regions of interest, the left and right DLPFCs. **B**. Averaged spectrograms (20 – 70 Hz) for each selected electrode (F4, F5, and F6 for the right DLPFC, and F3, F5, and F7 for the left DLPFC), in the 1 second 3000-4000ms window of the delay epoch.

### 3.3 Total Gamma Power

Total gamma power was derived from the 3000-4000ms of the delay period and from the 1 second inter-trial fixation epoch using the FFT from the left and right DLPFC. We found that power significantly decreased with age (*F* = 4.76, *p* = 0.03; Figure 4). Age by epoch (delay and fixation) interactions were not significant (p value), thus both delay and fixation epochs were included in the model and epoch was modelled as a covariate.

**Figure 4.**
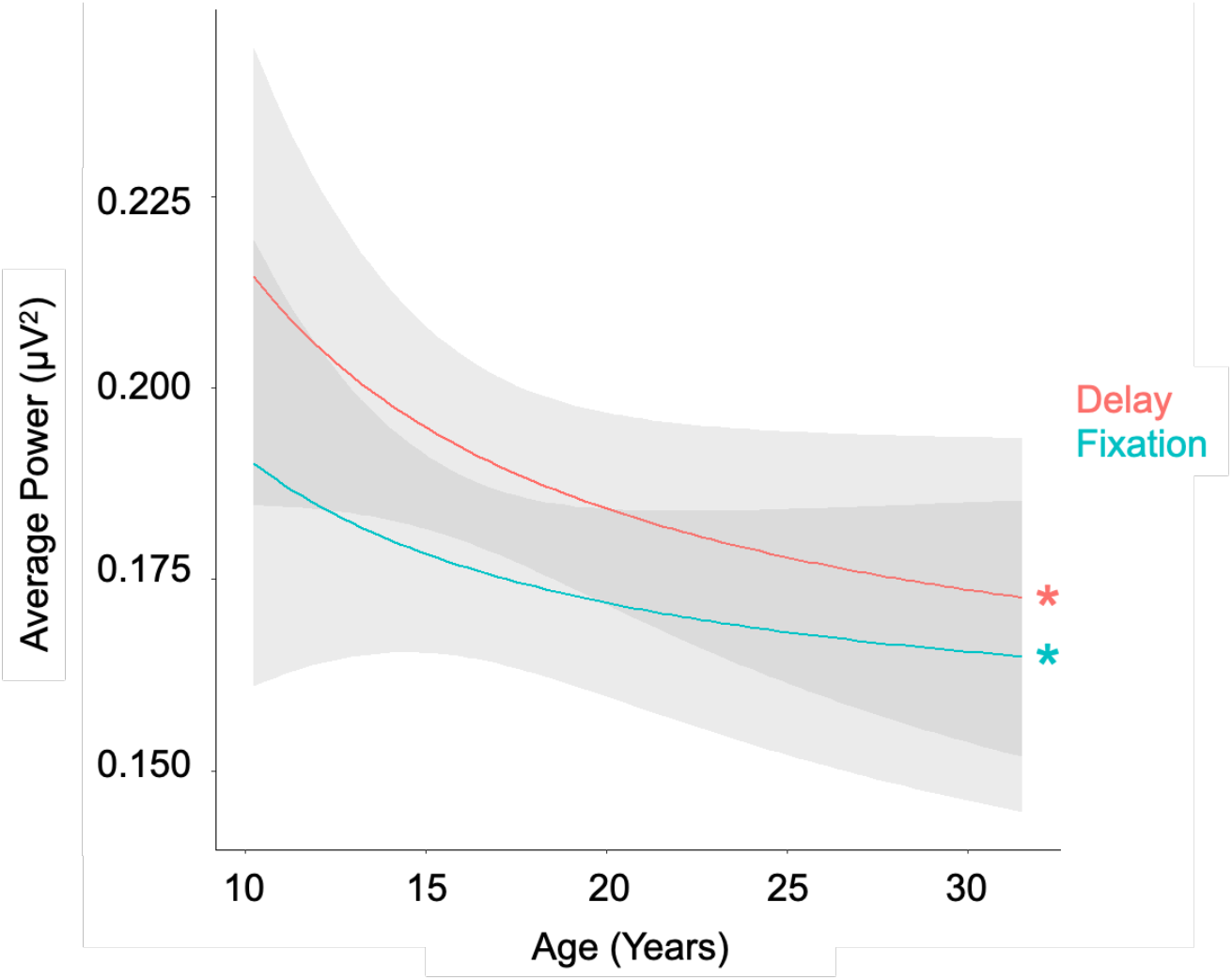
Average gamma power averaged across all trials. (* p < 0.01)

### 3.4 Gamma Emerges as Transient Events on Individual Trials

Based on evidence indicating that gamma bursts support maintenance of information in working memory^23,28,30^, we were interested in characterizing age-effects through adolescence in gamma events at the single trial level. Due to non-negative power spectral values, average power across trials can create a continuous band of activity, appearing as a sustained rhythm, even if the trial level response is better described by a series of discrete gamma events. To assess this in our data, we first visualized trial-by-trial gamma events during the same 3000-4000ms time window of the delay epoch used in the total power analysis above (see representative trials shown in Fig 5). These results demonstrate a pattern in which gamma band activity emerges from a series of transient increases in power (i.e., events) on individual trials, occurring at different moments within the epoch across trials. The presence of event-like gamma activity in the trial level spectrograms suggests the temporal dynamics of these events may be critical to understanding their contributions to working memory^27^, and provide a more mechanistic description of developmental change.

**Figure 5.**
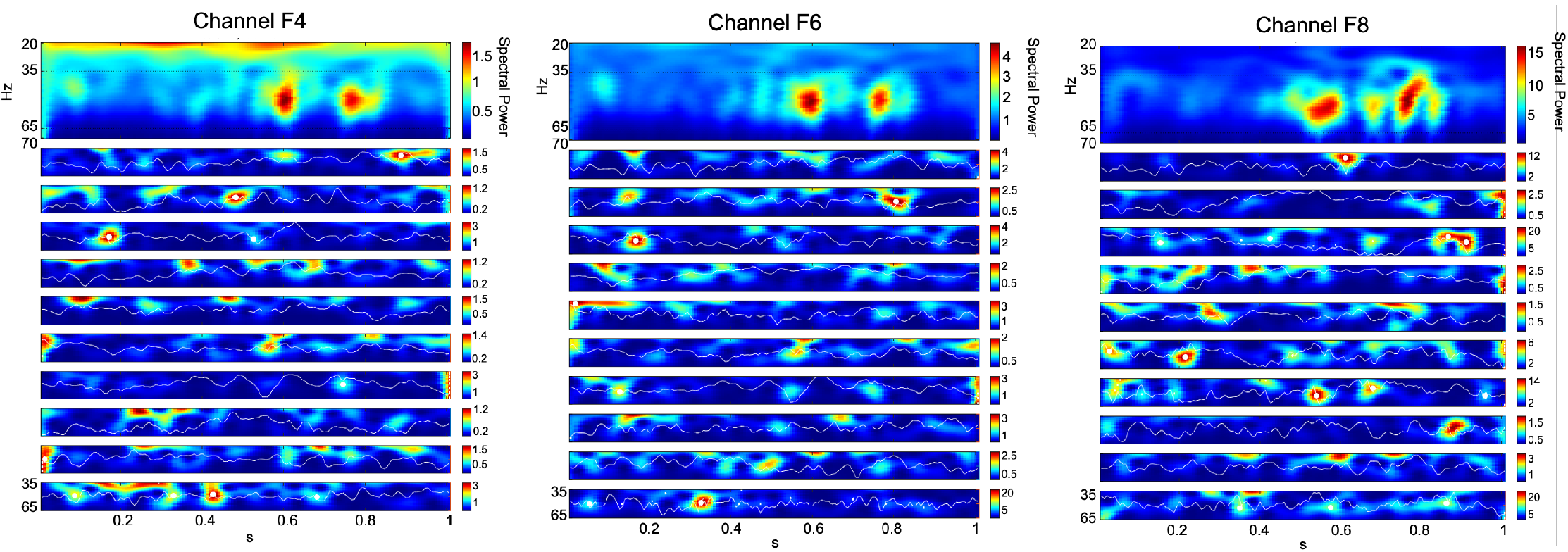
Gamma emerges as transient events in non-averaged spectrograms. Top panel shows averaged spectrogram (20-70 Hz) in the 1s 3000-4000ms window during the delay period, from a representative subject for the F4, F6, and F8 electrodes. The bottom panels show examples of delay period gamma band (35-65 Hz) activity in 10 randomly selected trials from the same participant, for the same electrodes. White dots denote local maxima in the spectrogram with maxima power above 6X median power of the maxima frequency.

### 3.5 Developmental Differences in EEG Spectral Event Measures

Spectral measures were computed across all trials for all participants spanning the full age range of our sample (10-32yo). Mean gamma event power significantly decreased across adolescence across both hemispheres (*F =* 51.5, *p* = 5.71e-06; Figure 6A), and we also observed a significant interaction of age with hemisphere (F = 1.72, *p* = 0.017; Figure 6A), indicating a greater decrease in the left DLPFC, and a significant interaction of age with epoch (F = 6.9, *p* = 0.0013; Figure 6A), indicating a greater decrease in the fixation epoch. Gamma event power variability showed decreases across age (F = 59.97, *p* = 1.2e-15; Figure 6D) and a significant interaction of age with epoch (F = 2.51, *p* = 0.0096; Figure 6D), suggesting a greater decrease in the fixation epoch. We did not observe a significant main effect of age on event number (F = 12.9, *p =* 0.22; Figure 6B) but did observe a main effect of epoch (β = 0.09, t = 5.8, *p* = 1.94e-08; Figure 6B), and a significant interaction of age with epoch (F = 3.370, *p* = 0.004; Figure 6B), indicating a greater decrease in the fixation epoch. Figure 7 illustrates the epoch by age interaction, showing the developmental differences between the number of events in the fixation epoch relative to the number of events in the delay period. We observed significant main effects of hemisphere (β= 0.07, t = 3.4, *p* = 0.027; Figure 6E) and epoch (β = 0.11, t = 7.4, *p* = 1.22e-08; Figure 6E) on event number variability, but no effects of age (all *p* > .05). There was a main effect of epoch on event duration (β = -0.0005, t = -3.4, *p* = 0.0093; Figure 6C), but no significant effects of age (F = 1.68, *p* > 0.05; Figure 6C) or age by epoch (F = 2.55, *p* = 0.65; Figure 6C) nor hemisphere (F = 0.42, *p* > 0.05; Figure 6C). Finally, event duration variability increased with age (F = 36.89, *p* = 0.00033; Figure 6F), had a main effect of region (β = 0.00044, t = 4.3, *p* = 0.025; Figure 6F), and a significant interaction of age with region (F = 3.81, *p* = 0.011; Figure 6F), indicating a significant increase in the left DLPFC.

**Figure 6.**
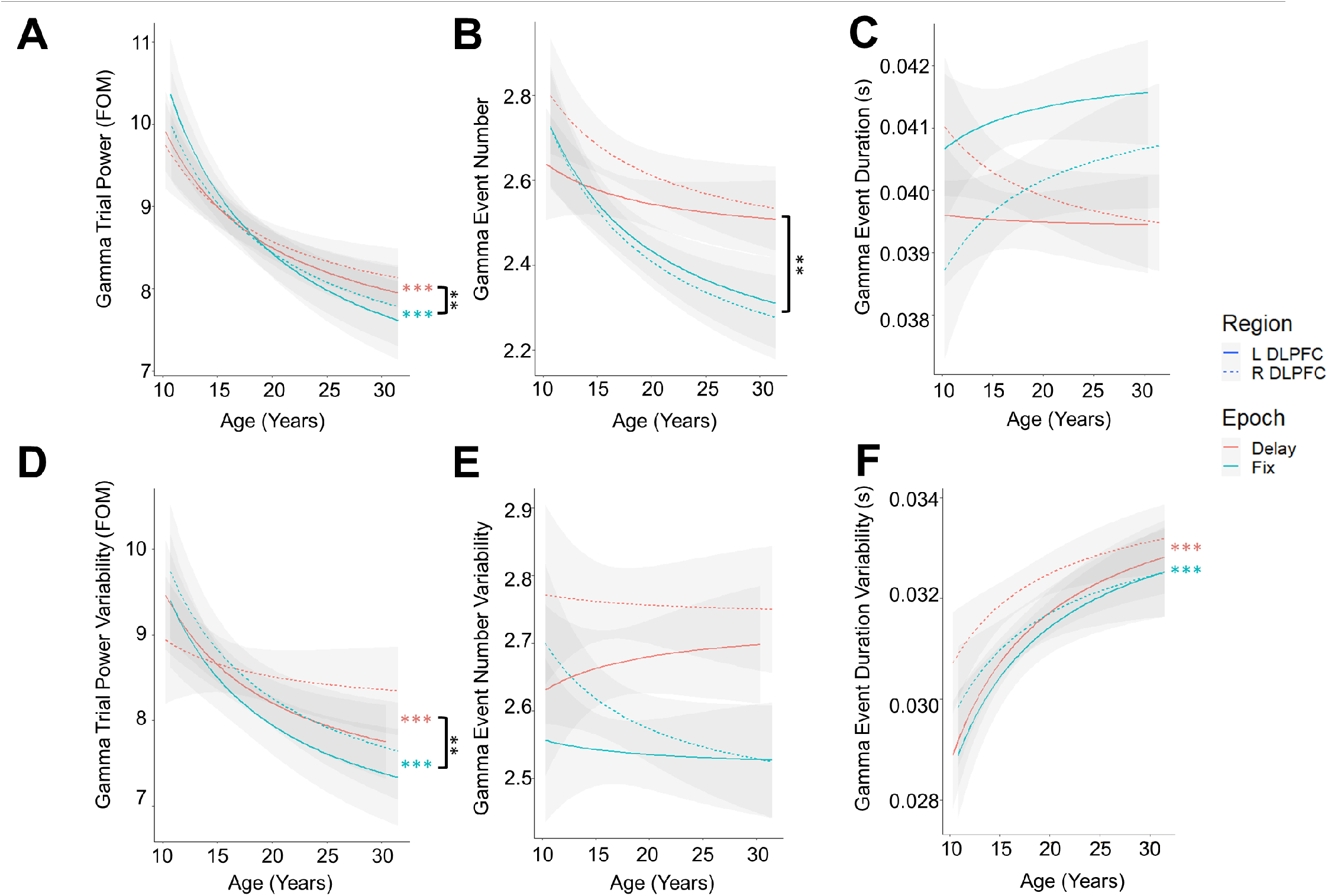
MGS delay (red) and fixation (blue) period EEG spectral measures in the left (solid line) and right (dotted line) DLPFCs. **A**. Mean trial power of a spectral burst. **B**. Mean number of spectral bursts per trial. **C**. Mean duration of spectral bursts per trial. **D**. Trial-by-trial variability of spectral burst power. **E**. Trial-by-trial variability of number of spectral events. Black lines and asterisks denote a significant epoch by age interaction. (* p < 0.01; ** p < 0.001; *** p < 0.0001)

**Figure 7.**
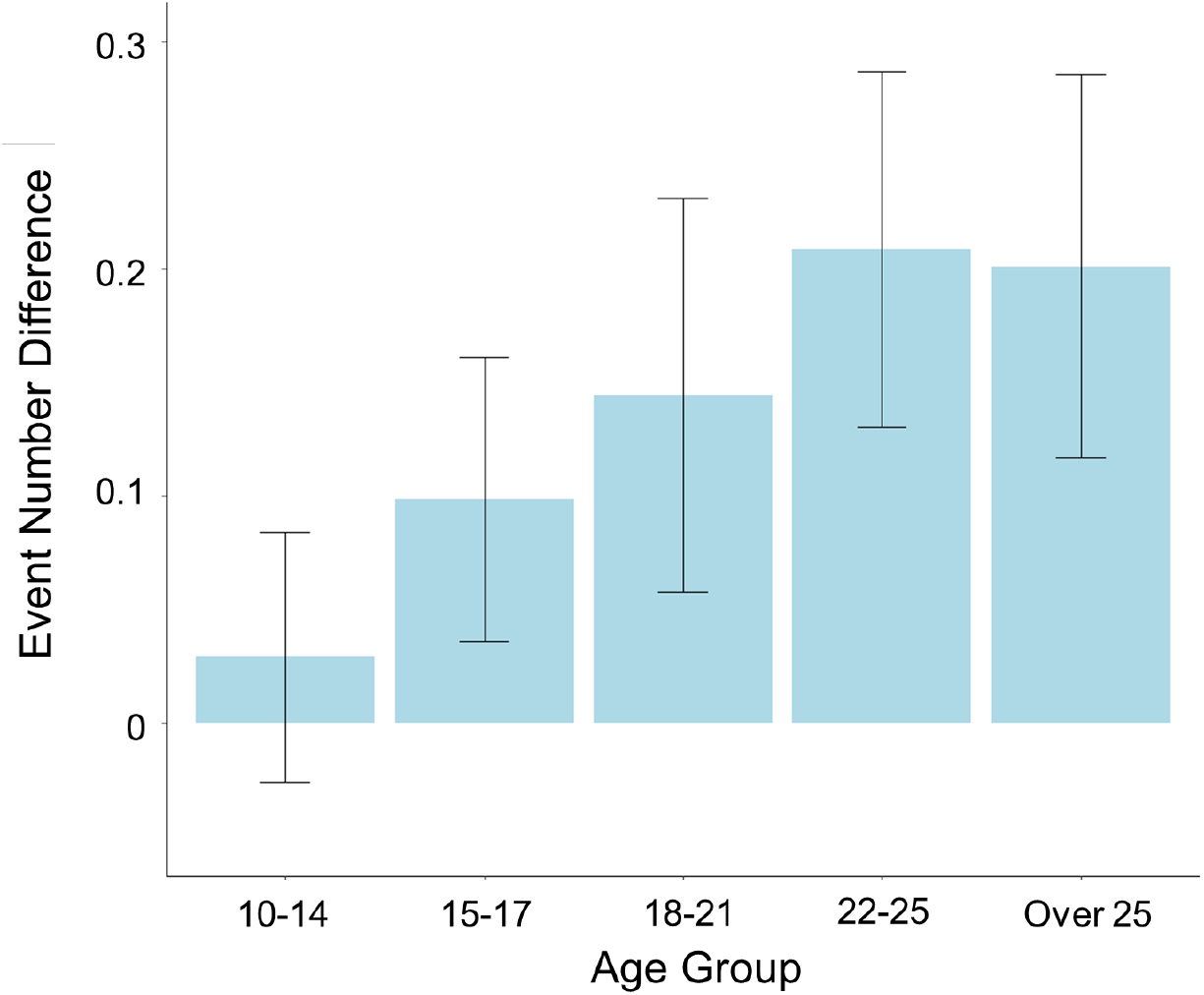
Gamma Event Number Difference between Delay and Fixation Epochs. The average number of gamma events for each age group from the fixation epoch was subtracted from the delay period. Error bars represent the standard error.

### 3.6 EEG Spectral Events Association with Behavior

We found a trend-level association between trial power and MGS accuracy; whereby somewhat higher trial power was associated with poorer MGS accuracy (increase in degrees from target; *p* = 0.057; Figure S1A). There were no significant associations between trial power variability and accuracy variability, nor event number and accuracy. Increased event number variability showed trends with higher accuracy variability (*p* = 0.06; Figure S1E) while controlling for hemisphere. Further statistics can be found in supplement table 1.

No associations between trial power and response latency nor trial power variability and response latency variability were found. No associations between event number and latency nor event number variability and response latency variability were found. Event duration and response latency did not have a significant relation (*p* = 0.11; Figure S2C). Hemisphere had a main effect on event duration variability and response latency variability (*p* = 0.032; Figure S2F). Further statistics can be found in supplement table 1.

### 3.7 Spectral Event Power Partially Mediates Total Power

We conducted mediation analyses to test whether the spectral event measurements exhibiting significant agerelated change (i.e., power and number of events) could account for age-related changes seen in the total gamma power measures in both the delay and fixation periods of the MGS task, similar to what has been previously reported in the literature.

Mediation analyses tested whether the relationship between age and total power was mediated by each spectral measure to determine the contribution of the spectral event measures to total gamma power. The effect of age on total power was partially mediated via event power in both the delay (Figure 8; β = 0.001, 95 % CI [0.0002, 0], *p* < 0.01) and fixation (Figure 8; β = 0.0017, 95 % CI [0.0007, 0], *p* < 0.0001) epochs. However, the number of gamma events did not mediate age-related changes in total power in either the delay (Figure 8; β = 0.0002, 95 % CI [-0.0001, 0], *p* > 0.05) nor fixation (Figure 8; β = 0.0005, 95 % CI [-0.0002, 0], *p* > 0.05) epochs.

**Figure 8.**
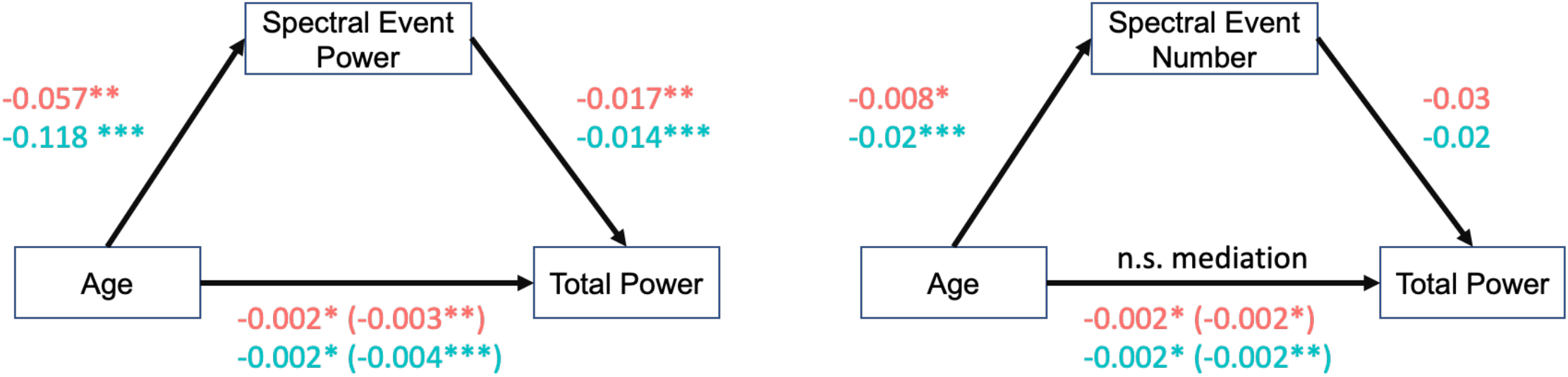
Left: mediation analysis to investigate whether spectral event power partially accounts for the agerelated changes in gamma band total power in the delay (red) and fixation (blue) epochs. Right: mediation analysis to investigate whether the number of spectral events partially accounts for the age-related changes in gamma band total power in the delay (red) and fixation (blue) epochs.

## 4. DISCUSSION

In this study we aimed to investigate the neural activity underlying working memory maintenance across adolescence. Consistent with previous literature^1,2^, working memory performance improved into adulthood including increased accuracy and decreased response latency to make a goal-directed response. We focused on understanding age-related differences in trial level transient events of activity during the delay period of a working memory task^23,28,30,31,43,44^, as opposed to sustained activity, as recent evidence has suggested its importance for understanding mechanisms underlying working memory delay period dynamics^21,27,28,45,46^. It is traditionally understood that sustained neural activity during working memory delay periods underlies the retention of the information in working memory. However, newer non-human primate evidence suggests that the sustained nature of this signal may arise due to the averaging of delay period activity across trials, while trial-level activity reveals transient neural events^21^ which support working memory maintenance. In this study, we leveraged the direct assessment of neural processing provided by scalp EEG, primarily reflecting postsynaptic cortical pyramidal cell activity^47,48^, with high temporal resolution that can characterize signal variability across timescales and frequency bands. Specifically, we investigated age-related effects from childhood to adulthood in the spectral events within the gamma frequency band, which has been found to support working memory maintenance^23,28^, and investigated their associations with working memory maintenance, and assessed whether transient events of activity at the trial level underlie age-related change in sustained activity and could thus further inform the underlying mechanisms of the development of working memory through adolescence.

To determine regions of interest for investigating age-related differences in transient gamma events during the working memory delay and fixation epochs, average gamma power was calculated across our adult cohort and plotted across brain topography. Right and left DLPFC regions of interest were selected, both for the presence of gamma band activity as seen in Figure 3, and for the previously reported involvement of the DLPFC in working memory tasks^22,49–53^. Our results show that total gamma power, derived from the fast Fourier transform (FFT), decreases through adolescence into young adulthood (Figure 4), consistent with previous studies of gamma band activity^15^. Such age-related decreases have also been reported in the beta^12^ and theta^14^ bands, suggesting that decreases in oscillatory power may be a common developmental pattern. Interestingly, we found that on individual trials gamma band activity was present in discrete, brief events, and only when averaging across trials did more sustained activity patterns emerge (Figure 5). This finding is consistent with previous studies that have shown that beta and gamma events occur during encoding and then reappear during the delay period of working memory tasks^28,30,31^. Expanding upon the developmental decreases seen in the gamma band across trials, we then investigated age-related change in the temporal dynamics of these transient gamma band spectral measures. We found that mean gamma event power, event power variability, and the duration variability of an event, all significantly decreased through adolescence, in both the delay period and fixation epochs of the working memory task (Figure 6). These results are in agreement with studies showing decreases in power through adolescence across the cortex^12,16,54–58^. Furthermore, previous human EEG studies have characterized elevated gamma frequency power in local field potentials during working memory maintenance^59–62^, that decreases across adolescence, suggesting optimization of encoding and maintenance circuitry^19^. Variability in EEG signals has been found to decrease in parallel to developmental decreases in behavioral variability^63^, and stabilization of neural signaling occurs in parallel to maturation of structural^64^ and functional ^65–69^ connectivity. Thus, decreases in variability may reflect stabilization of neural function and behavior. The developmental changes seen in both the delay and fixation epochs suggest that basic dynamics of gamma oscillations specialize through adolescence regardless of cognitive demands. Myelination and synaptic pruning across association cortex at this time, and in prefrontal cortex in particular^12,15,45,57,70^, may underlie optimization of information processing resulting in decreased need to engage mechanisms supported by gamma oscillations into adulthood. Importantly, gamma event power significantly mediated the changes we identified in total gamma band power throughout the fixation and delay period epochs, suggesting that this may provide a mechanistically specific driver of these frequently reported developmental changes in gamma band power.

Interestingly, we also identified developmental change in the number of gamma events in both the delay and fixation epochs. However, unlike event power and duration, the number of gamma events (i.e., the number of events per second) exhibited a significant age by epoch interaction, reflecting that the number of gamma events in the fixation period of the working memory task decreased more drastically with age than during the delay period. These results may suggest that there is a baseline decrease in the number of gamma events with age, as indicated by the decrease of events in the fixation epoch, possibly due to synaptic pruning^10,12,15,57,71^ or reduction in spontaneous activity^2,45^. In contrast, the number of gamma events during the working memory delay period did not appear to decrease. These findings resulted in age-related increases in the difference between number of events supporting fixation compared to working memory maintenance so that by adulthood there are a greater number of events supporting working memory maintenance compared to fixation rest (Figure 7). This finding suggests that with development gamma events become specialized for working memory maintenance and may support working memory development. Previous work has shown the presence of gamma events during working memory decoding^28^, ramping up in anticipation of WM readout^23^, and have suggested events represent attractor states that correspond to working memory information^28^. Importantly, in addition to showing task-dependence and epoch specificity, which was not observed for event power or duration, the number of gamma events was not strongly related to total gamma power as indicated by the non-significant mediation for both the fixation and delay period epochs. Together, these results suggest that the rate of gamma events may represent a developmental feature of gamma band processing that is distinct from previously reported changes in gamma power. Thus, our results support that the relative increase in gamma events specific to the working memory maintenance period may thus be an important and novel component of developmental refinements in working memory performance. However, a limitation of our study is that while these changes in gamma events were specific to the working memory maintenance period, they did not explain individual differences in performance, and thus further work is necessary to clarify their specific behavioral contribution.

Basic working memory function is available in infancy^72^, and at the trial level, adult level performance can be attained even in childhood, though greater variability across trials results in decreased mean performance^5^. Thus, these results suggest that the core mechanisms of working memory are already available in adolescence. Variability in EEG signals has been found to decrease in parallel to developmental decreases in behavioral variability^63^. Stabilization of neural signaling occurs in parallel to maturation of structural^64^ and functional ^65–69^ connectivity. Thus, decreases in variability may reflect stabilization of neural function and behavior. Variability in gamma events during maturation in adolescence may serve as an adaptive exploratory approach^5^, which by adulthood has already reached optimal performance and may be more related to variability in attention. The results observed here in adolescence are in accord with neurophysiological data showing that saccade reaction times are strongly correlated with firing activity^73–75^, with higher rates of neuronal firing being linked to faster behavioral responses^73,75^. Together, these results suggest that from adolescence to adulthood there may be a process of specialization in which greater neural function may support best performance in adolescence, followed by a shift to less neural activity being required for adult level optimal executive function.

We focused on understanding age-related differences in trial level activity as recent evidence indicates its importance for understanding mechanisms underlying working memory delay period dynamics^21,27,28,45,46^. Thus, we sought to investigate whether trial level transient events of activity underlined the traditionally observed sustained activity. Our mediation analyses suggested as much, with trial level transient event power partially mediating the age-related changes in total sustained power in both the delay and fixation epochs. However, the number of gamma events did not significantly mediate total sustained power. These results suggest that while the spectral event measures are related to trial averaged power, they may be driven by different neural mechanisms^76^ supporting unique aspects of executive function. Recent work shows compelling evidence that transient events during delay periods support maintenance of mnemonic information through the delay period^31^. More specifically, previous work has shown that working memory activity is not stationary, and it has been hypothesized that information is conveyed as spiking in short attractor states and held by synaptic changes in-between states^28^. Our findings that trial level gamma activity decreases with development provides compelling new evidence for mechanistic changes in neural processing characterized by refinements in neural function as behavior becomes optimized in adulthood.

This study is limited by a cross-sectional cohort, which impede inferences regarding within-subject developmental effects, and future studies leveraging longitudinal cohorts will be necessary to ascertain whether age-related changes in EEG measures support developmental improvements in working memory performance. Further, while we did not find associations between spectral events measures and performance, this may not be surprising as working memory involves a wide circuitry of neuronal activity. Future work utilizing a working memory task with variable load conditions may provide further insight into the behavioral contributions of gamma events since gamma oscillations have been found to be correlated with working memory load^60,77,78^.

Together, these results suggest that critical refinements in neural function underlie improvements in working memory from adolescence to adulthood, reflecting the ability to readily and consistently engage mechanisms to employ existing cognitive processes, that during adolescence may leverage increases in neural dynamics and variability to improve performance at the trial level. Characterizing neural mechanisms that underlie normative development can inform impaired trajectories, including psychiatric disorders, where working memory is predominantly affected such as in psychosis^79,80^ and depression^81,82^, which emerge during the adolescent period^83^.

## Acknowledgements

We thank the University of Pittsburgh Clinical and Translational Science Institute (CTSI) for help in recruiting participants. We thank Matthew Missar and Alyssa Famalette for their work involving our data collection.

