## Supplement for "Age-related change in transient gamma band activity during working memory maintenance through adolescence"

**Supplementary Figures and Tables**

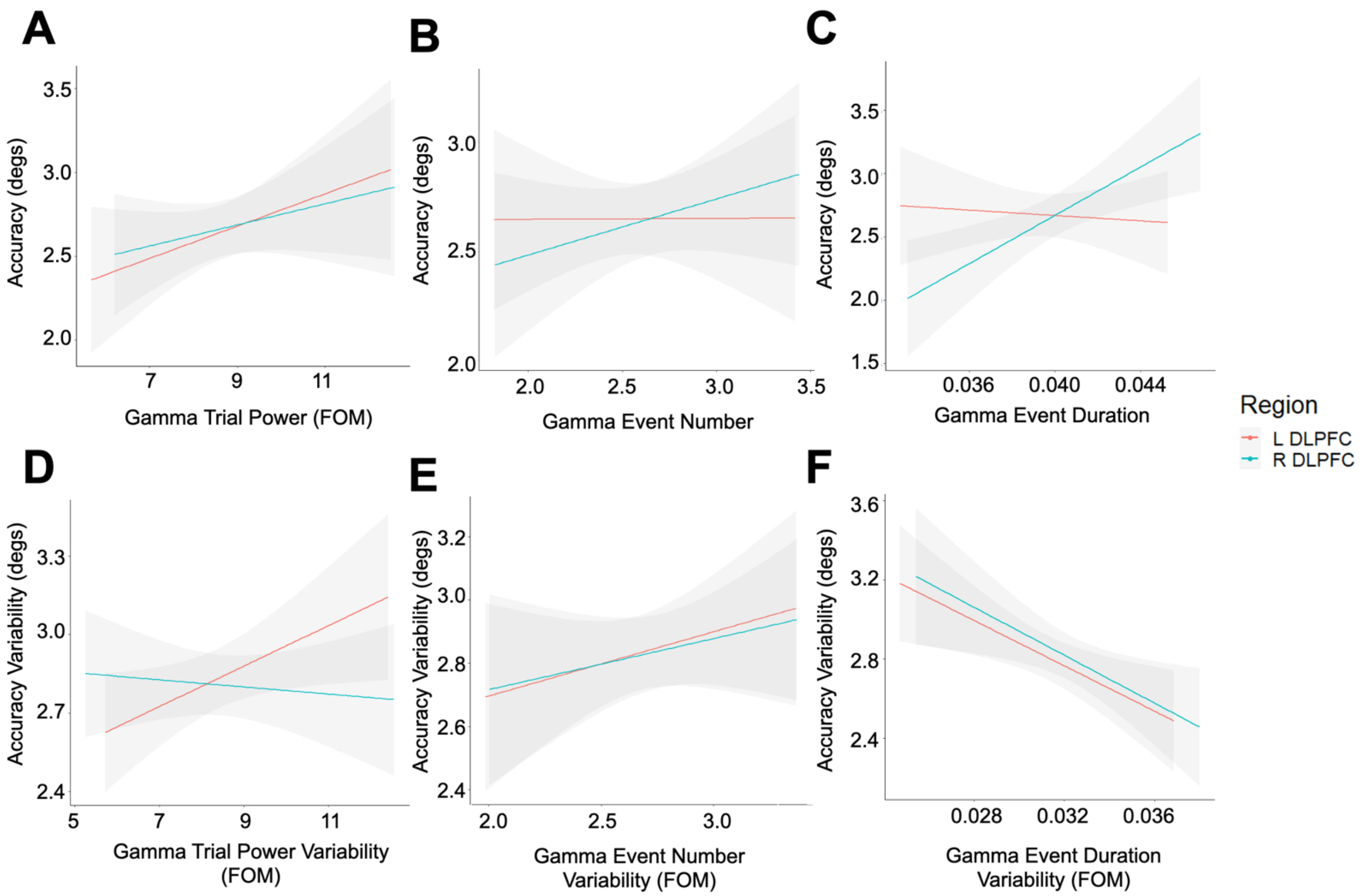

**Supplementary Figure S1**. MGS performance measures of accuracy (measured in degrees away from the target location) vs delay period EEG spectral measures. **A.** Accuracy vs gamma trial power for the left (red) and right (blue) DLPFC. **B.** Accuracy vs number of gamma events for the left (red) and right (blue) DLPFC. **C.** Accuracy vs duration of a gamma event for the left (red) and right (blue) DLPFC. **D.** Accuracy trial variability vs gamma trial power variability for the left (red) and right (blue) DLPFC. **E.** Accuracy trial variability vs gamma event number variability for the left (red) and right (blue) DLPFC. **F.** Accuracy trial variability vs gamma event duration variability for the left (red) and right (blue) DLPFC.

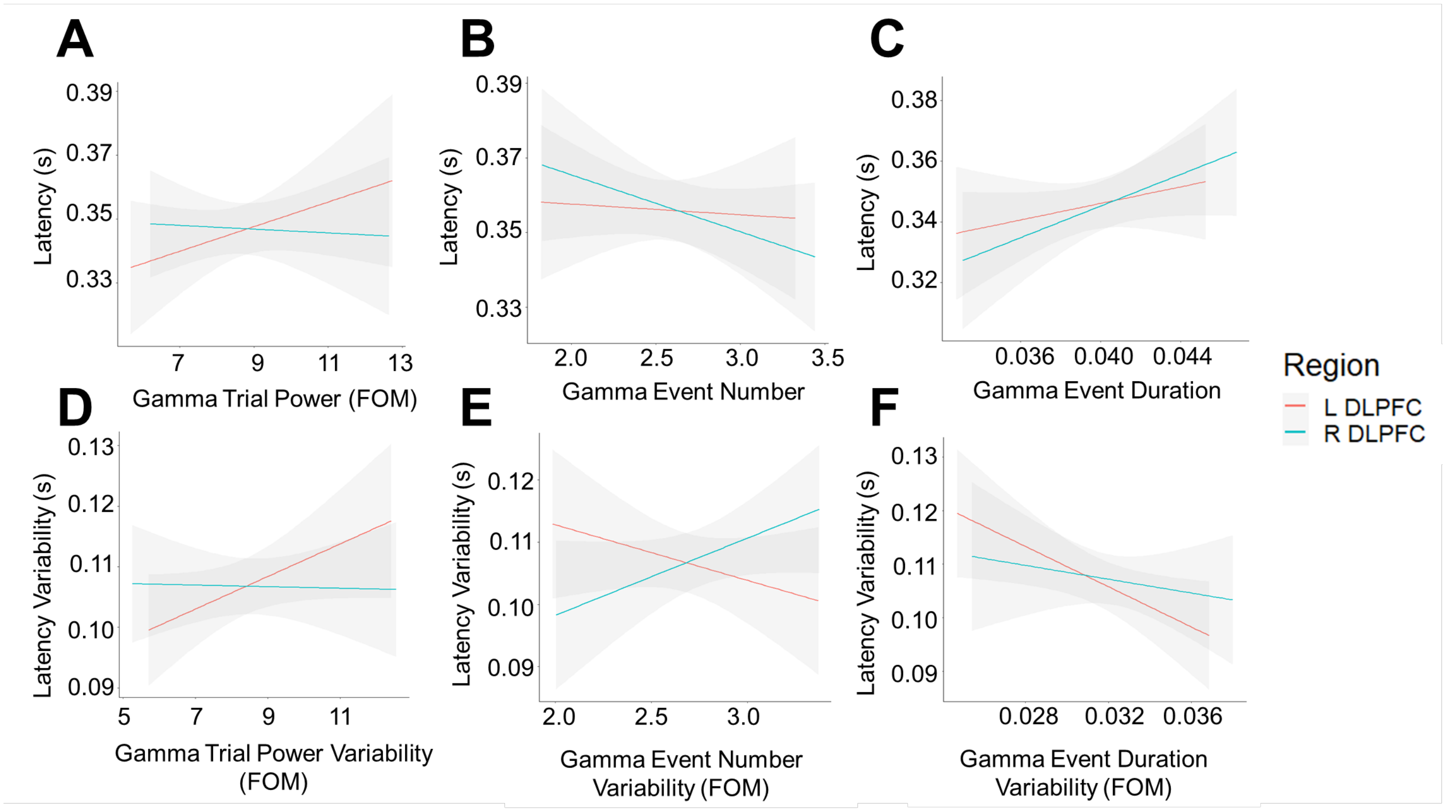

**Supplementary Figure S2**. MGS performance measures of response latency and vs delay period EEG spectral measures. **A.** Latency vs gamma trial power for the left (red) and right (blue) DLPFC. **B.** Latency vs number of gamma events for the left (red) and right (blue) DLPFC. **C.** Latency vs duration of a gamma event for the left (red) and right (blue) DLPFC. **D.** Latency trial variability vs gamma trial power variability for the left (red) and right (blue) DLPFC. **E.** Latency trial variability vs gamma event number variability for the left (red) and right (blue) DLPFC. **F.** Latency trial variability vs gamma event duration variability for the left (red) and right (blue) DLPFC.

**Supplementary Table 1**. Table of all Results. Top third: spectral events by age with epoch and hemisphere main effects and interactions with age. Middle third: spectral events by accuracy with age, epoch, and hemisphere main effects and interactions with accuracy. Bottom third: spectral events by latency with age, epoch, and hemisphere main effects and interactions with latency. (* $p < 0.01$; ** $p < 0.001$; *** $p < 0.0001$)

| | **Event Power** | **Event Number** | **Event Duration** | **Power Variability** | **Event Number Variability** | **Event Duration Variability** |
|---|---|---|---|---|---|---|
| Age | 5.71e-06 *** | n.s | n.s | 1.2e-15 *** | n.s | 0.00033 *** |
| **Covariates** | | | | | | |
| Epoch | n.s | 1.94e-08 *** | 0.0093 ** | n.s | 1.22e-08 *** | n.s |
| Hemisphere | n.s | n.s | n.s | n.s | 0.027 * | 0.025 * |
| Epoch:Age | 0.0013 ** | 0.004 ** | n.s | 0.0096 ** | n.s | n.s |
| Hemisphere:Age | 0.017 * | n.s | n.s | n.s | n.s | 0.011 |
| Accuracy | 0.057 | n.s | 0.26 | n/a | n/a | n/a |
| Accuracy Variability | n/a | n/a | n/a | n.s | 0.06 | n.s |
| **Covariates** | | | | | | |
| Epoch | n.s | n.s | n.s | n.s | n.s | n.s |
| Hemisphere | n.s | n.s | n.s | n.s | 0.018 * | 0.0055 ** |
| Epoch:Accuracy | n.s | n.s | n.s | n.s | n.s | n.s |
| Hemisphere:Accuracy | n.s | n.s | 0.025 | n.s | n.s | n.s |
| Latency | 0.25 | 0.198 | 0.11 | n/a | n/a | n/a |
| Latency Variability | n/a | n/a | n/a | n.s | n.s | n.s |
| **Covariates** | | | | | | |
| Epoch | n.s | n.s | n.s | n.s | n.s | n.s |
| Hemisphere | n.s | n.s | n.s | n.s | n.s | 0.032 * |
| Epoch:Latency | n.s | n.s | n.s | n.s | n.s | n.s |
| Hemisphere:Latency | n.s | n.s | n.s | n.s | n.s | n.s |
